# No cardiac phase bias for threat perception under naturalistic conditions in immersive virtual reality

**DOI:** 10.1101/2024.01.31.578172

**Authors:** Felix Klotzsche, Paweł Motyka, Aleksander Molak, Václav Sahula, Barbora Darmová, Conor Byrnes, Iveta Fajnerová, Michael Gaebler

**Author notes:** Corresponding authors: Felix Klotzsche, Paweł Motyka. These authors contributed equally (Felix Klotzsche, Paweł Motyka).

## Abstract

Previous studies have found that threatening stimuli are more readily perceived and more intensely experienced when presented during cardiac systole compared to diastole. Also, threatening stimuli are judged as physically closer than neutral ones. In a pre-registered study, we tested these effects and their interaction using a naturalistic (interactive, 3D) experimental design in immersive virtual reality: We briefly displayed threatening and non-threatening animals (four each) at varying distances (1.5–5.5 meters) to a group of young, healthy participants (n = 41), while recording their ECGs (electrocardiograms). Participants then pointed to the location where they had seen the animal (ca. 29k trials in total). Our pre-registered analyses indicated that perceived distances to both threatening and non-threatening animals did not differ significantly between cardiac phases – with Bayesian analysis supporting the null hypothesis. There was also no evidence for an association between subjective fear and perceived proximity to threatening animals. These results contrast with previous findings that used verbal or declarative distance measures in less naturalistic experimental conditions. Furthermore, our findings suggest that the cardiac phase-related variation in threat processing may not generalize across different paradigms and may be less relevant in naturalistic scenarios than under more abstract experimental conditions.

**Impact statement:** To our knowledge, this is the first study to assess the influence of interoceptive cardiac signals on visual perception using naturalistic stimuli in immersive virtual reality. We based the design of our experiment on previous reports about threat processing biases and the role of the cardiac cycle but did not observe the expected effects. This poses the question of their generalizability from abstract settings to more naturalistic setups, featuring greater behavioral engagement and a richer sensory environment.

## Introduction

Detecting threats and appropriately reacting to them supports an organism’s physical integrity and survival. Perceiving or imagining a (potential) danger typically results in the feeling of fear. The standard fear response mobilizes psychological and physiological resources to effectively face or avoid the threat (“fight-or-flight”; Cannon, 1915). While exposure to fearful stimuli is known to influence the heart rate (Hare, 1973; Mocaiber et al., 2011; Palomba et al., 2000; Ruiz-Padial et al., 2011), cardiac activity, in turn, also affects fear processing (Garfinkel et al., 2021; Garfinkel & Critchley, 2016): When presented at cardiac systole (i.e., when the heart muscles contract to eject the blood into the arteries), fearful faces are more easily detected (Garfinkel et al., 2014; but see also Leganes-Fonteneau et al., 2021) and rated as more intense (Garfinkel et al., 2014; Leganes-Fonteneau et al., 2021) than when presented at cardiac diastole (i.e., when the heart muscles relax and the heart refills with blood). Also, the expression of threat-related stereotypes (Azevedo et al., 2017) and attention to fearful stimuli (Azevedo et al., 2018) were shown to be more pronounced during cardiac systole. The modulatory effects of the cardiac cycle on perception have also been reported for non-emotional stimuli in the visual (Réquin & Brouchon, 1964; Sandman et al., 1977; but see also Elliott & Graf, 1972) the somatosensory (Al et al., 2020; Motyka et al., 2019; Wilkinson et al., 2013), and the acoustic domain (Schulz et al., 2009). Apart from perception, action also varies across the cardiac cycle: Eye movements (Galvez-Pol et al., 2020; Ohl et al., 2016) and button presses (Kunzendorf et al., 2019) are increased during systole, while movements related to tactile exploration have longer durations when initiated during systole (Galvez-Pol et al., 2022).

Such effects of cardiac timing on perception and action could be physiological artifacts – non-functional byproducts of our body’s normal functioning. For example, heartbeat-related shifts in blood flow and blood pressure can introduce “noise” that interferes with activity in sensory organs (e.g., the retina; Joseph et al., 2019; Tornow et al., 2018), peripheral organs responsible for action (e.g., muscles; Birznieks et al., 2012; Fairfax et al., 2013), and the brain (Elbert & Rau, 1995; Rau et al., 1993; Rau & Elbert, 2001). Such periodic visceral signals may also be predicted and integrated into (active) perceptual processing (e.g., in the posterior insula; Hsueh et al., 2023; Klein et al., 2021), among other things, to counter detrimental effects, such as sensory attenuation or uncertainty (Al et al., 2020; Allen et al., 2022). In this view, cardiac timing effects may be behaviorally relevant and evolutionarily adaptive – at least for the processing of specific types, such as threat-related or other motivationally relevant stimuli (Garfinkel & Critchley, 2016; Pramme et al., 2016; Schulz et al., 2020). However, the extent to which cardiac-cycle biases hold in real-world or everyday-life situations remains unclear as empirical evidence beyond artificial, decontextualized laboratory experiments is missing.

Immersive virtual reality (VR) technology facilitates more naturalistic (i.e., dynamic, interactive, and less decontextualized) neuroscientific studies by completely surrounding the observer with interactive, computer-generated scenarios that are contextually rich (Diemer et al., 2015). As the virtual environment is artificially and purposefully created, a high level of experimental control can be maintained. Such more naturalistic experiments allow to study the organism under conditions it was optimized for (Gibson, 1979; Hasson et al., 2020), and their findings may more readily generalize to real-world circumstances and provide better models of mind-brain-body functioning (Matusz et al., 2019; Shamay-Tsoory & Mendelsohn, 2019). In the present study, we employed a naturalistic VR task to gauge the real-world relevance of cardiac phase biases in the visual perception of threatening and non-threatening objects.

One of the perceptual biases supporting adaptive behavioral responses to threats is the underestimation of distance to threatening stimuli. Such an effect has been demonstrated for fear-evoking animals and humans (Cole et al., 2013; Fini et al., 2018), as well as for an initially neutral stimulus that became associated with pain (Tabor et al., 2015). Threatening animals are also perceived as approaching more quickly than non-threatening ones (Basanovic et al., 2019; Vagnoni et al., 2012; Witt & Sugovic, 2013). These kinds of amplification of threat perception are believed to facilitate faster responses in the face of danger (Balcetis & Cole, 2014; de Carvalho, 2022). To the best of our knowledge, no studies have yet examined whether this proximity bias for threats is linked to cardiovascular fluctuations and, more generally, whether these fluctuations can affect the spatial representation of threatening objects. Notably also, previous studies investigating proximity bias predominantly relied on verbal estimations (Cole et al., 2013; Tabor et al., 2015), which, in general, are considered less accurate than behavioral measures (e.g., Andre & Rogers, 2006; Etchemendy et al., 2018; Kunz et al., 2009) and more susceptible to demand characteristics such as the study setting or experimenter influences (Firestone & Scholl, 2016). We suggest that stereoscopic VR is particularly suited to test functional distance perception and estimation. It features a more naturalistic depth perception than 2D screens (as often used in classical experiments, e.g., Kim & Harris, 2022; Tabor et al., 2015). Thereby it presents itself as a suitable platform for exploring influences on distance perception in a more naturalistic manner. This allows for the investigation of whether effects found in more abstract laboratory settings can also be observed in conditions that are closer to real-world situations. Additionally, when compared to field experiments or observational studies, it offers more clear-cut experimental control and precise spatial measurements.

We leveraged the advantages of immersive VR to investigate the effects of the cardiac cycle on perceived distances to threatening and non-threatening visual objects. We designed a novel task using immersive VR, in which participants behaviorally indicated the perceived position of realistic 3D animals. The stimuli were presented briefly at different phases of the cardiac cycle and at various distances from the observer. In this pre-registered study (https://osf.io/a7n9b/), we hypothesized that threatening stimuli are perceived as closer than non-threatening ones (Cole et al., 2013; Tabor et al., 2015). Based on the findings that the processing of threat-related signals is amplified during cardiac systole (Azevedo et al., 2017; Garfinkel et al., 2014, 2021; Leganes-Fonteneau et al., 2021), we also hypothesized that threatening stimuli are perceived as closer during earlier (i.e., systole) compared to later (i.e., diastole) phases of the cardiac cycle. Additionally, we explored associations between distance estimates and negative feelings (threat, disgust) evoked by the stimuli as well as individual anxiety levels.

## Method

### Participants

We acquired data from 46 healthy participants (28 females, mean age = 28.9 ± 4.6 years) with normal or corrected-to-normal vision, and without (self-reported) psychiatric, neurological or cardiovascular conditions. Due to technical issues, data from 5 participants were incomplete or corrupted, yielding a final sample of 41 participants (24 females, mean age = 28.8 ± 4.4 years, range: 19–39 years). The pre-registered target sample size (N = 40) was chosen to be similar to that used in previous cardiac-timing studies (Garfinkel et al., 2014; Kunzendorf et al., 2019; Ohl et al., 2016). Participants were recruited through posters at University buildings and the database of the Berlin School of Mind and Brain. They were informed that realistic 3D models of animals would be presented, so individuals with fear of animals (e.g., arachnophobia) could decide not to participate in the study. All participants gave written informed consent before taking part in the study, and they were financially compensated for their participation. The procedure was approved by the Ethics committee of the Psychology Department at the Humboldt-Universität zu Berlin.

### Stimuli

Eight 3D models of animals served as stimuli, 4 threatening and 4 non-threatening ones (Figure 1). They were selected based on the results of an online study, in which an independent set of participants (N = 94, 61 females, mean age = 29.01 ± 5.95 years, range: 18–65 years) rated pictures of fourteen 3D models of animals. The models were sourced from the Unity Asset Store (https://assetstore.unity.com/). For the online ratings, renderings of the same virtual scene as in the immersive main experiment containing one animal per trial were shown. For each animal, participants answered the following questions: 1) “Please rate how threatening the presented animal is to you”; 2) “Please rate how disgusting the presented animal is to you”; 3) “Please rate how fast the presented animal can move towards you”. All responses were given on a 7-point Likert-type scale (min: “not at all”, max: “very much”). Based on the ratings, we performed a median split and selected a total of 8 animals: 4 rated as more and 4 as less threatening (Figure S1). Importantly, animals between both groups were approximately matched with respect to their size. Of note, threat and disgust ratings, as well as threat and movement speed ratings, were positively correlated (*r* = .66 and *r* = .61, respectively).

**Figure 1.**
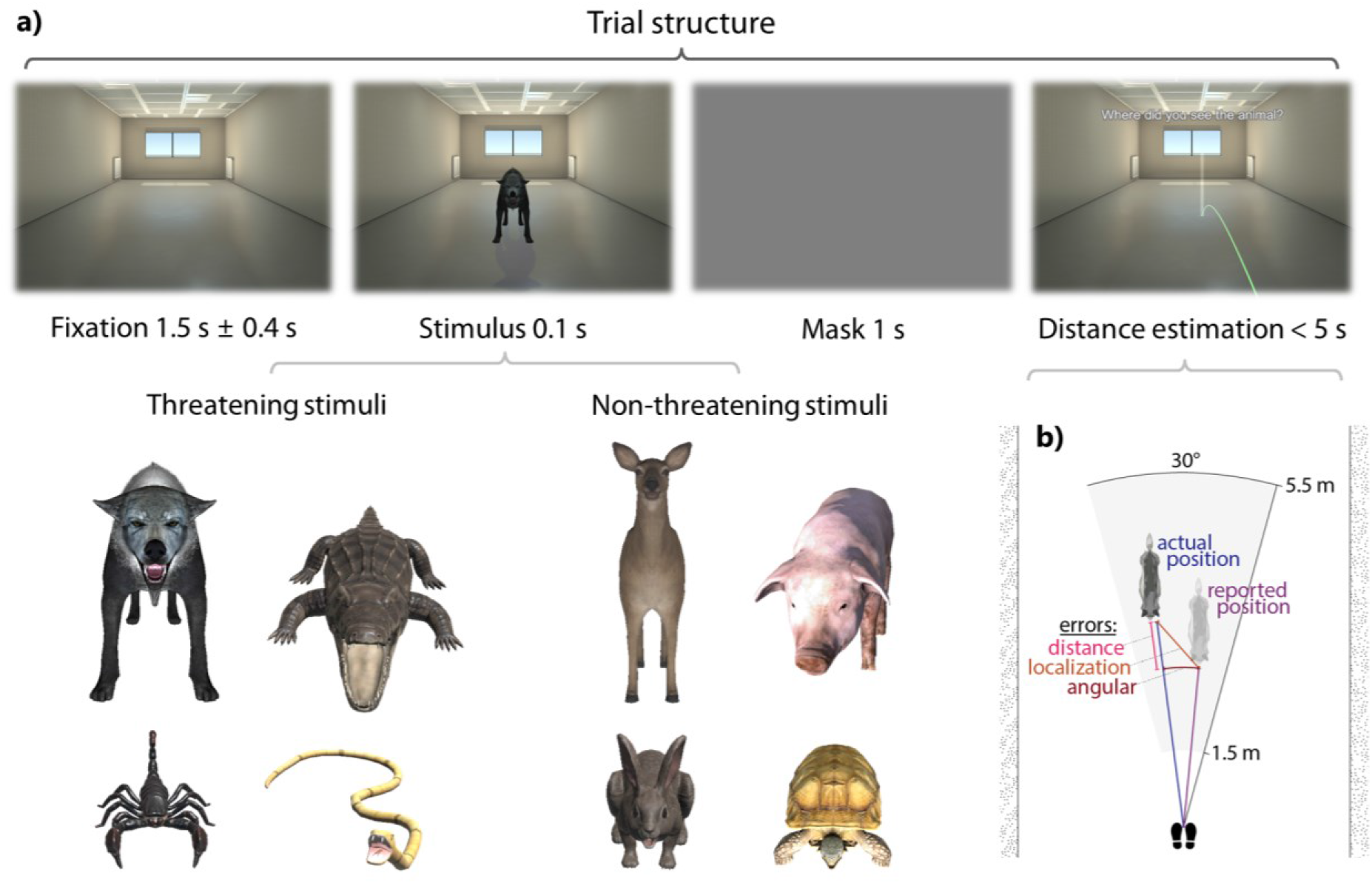
Experimental procedure. (**a**). Temporal structure of a single trial: After a fixation period, participants saw a threatening or non-threatening animal for 100 ms before the whole scene was masked and the empty room reappeared. Participants then indicated where they had perceived the closest point of the animal’s head (“the nose”). Animals were presented at varying distances (1.5–5.5 m) from observers, with the onset times being randomly distributed across the cardiac cycle. (**b**) The main dependent variable, referred to as "distance error," was the disparity between two distances: one from the participant to the reported animal location (purple line) and the other to its actual position (blue line). Negative distance errors signify underestimation, while positive errors indicate overestimation of the distance to the stimulus.

### Task and setup

At the beginning of each trial, participants were presented with an empty room for 1,500 ms ± 400 ms (jitter uniformly distributed). Next, a virtual animal, facing the observer, was displayed for 100 ms. The animal was positioned at a distance between 1.5 and 5.5 meters from the observer, with the exact distance randomly drawn from a continuous uniform distribution for each trial. We constrained the possible animal positions to a circular segment of 30° width centered around the participant and the long axis of the room (see Figure 1).

The stimulus presentation was immediately followed by the entire scene graying out for 1000 ms. Right after, participants used the handheld controller of the VR system to indicate the position in the (now empty) room at which they had perceived the closest point of the animal’s head. To do so, they moved a visual marker to the according location. The marker was a semi-transparent vertical bar (0.1 x 1.2 m; width x height) which could be moved along the floor plane of the virtual room by pointing with the VR controller. The implementation of an intuitive mapping between controller movement and the placement of the marker was based on the Teleport class in the SteamVR library (v2.0.1, Valve Corporation, Bellevue, United States). Apart from a virtual hand (glove) holding a digital replica of the VR controller, the body of the participant was not represented in the virtual environment.

During the main part of the experiment, participants’ electrocardiogram (ECG) was recorded (sampling rate: 1000 Hz; hardware-based lowpass filter at 262 Hz; third order sinc filter, –3 dB cutoff) via three electrodes connected to a LiveAmp EEG amplifier (BrainProducts GmbH, Gilching). Electrodes were attached according to a modified Einthoven procedure (Lead II; right clavicle, left hip bone, right ankle). Participants were seated and wore a VR head-mounted display (HTC Vive, HTC, Taiwan; refresh rate: 90 Hz). The VR environment was created using Unity (v2018.2.11f1; Unity Technologies, San Francisco, California). The virtual environment comprised an unfurnished, rectangular room measuring 5 by 15 meters. It included two windows and two radiators positioned at the opposite end from the participant who was located centrally in front of one of the short walls, facing towards the room (see Figure 1). The room provided perspective cues (linear perspective, foreshortening) and contained static pictorial depth cues (textured ceiling, light reflections on the floor) to aid distance perception (Renner et al., 2013). We provide an executable of the experimental VR software at https://osf.io/a7n9b/.

### Procedure

Upon arrival in the lab, participants completed a demographic questionnaire, the State-Trait Anxiety Inventory (STAI-S and STAI-T; Spielberger et al., 1970), and a shortened version of the Simulator Sickness Questionnaire (SSQ; Kennedy et al., 1993) comprising three items from the nausea (dizziness, nausea, general discomfort) and three items from the oculomotor subscale (headache, blurred vision, difficulty concentrating). This assessment, in combination with a second measurement of the SSQ at the end of the session, aimed to monitor changes in the participants’ condition before and after the VR experience, thereby determining if the experiment had any adverse effects (in line with previous recommendations; Bimberg et al., 2020; Brown et al., 2022; which depart from the original guidelines: Kennedy et al., 1993). To ensure that all participants had intact stereoscopic perception, they completed a Titmus test (Fly-S Stereo Acuity Test, Vision Assessment Corporation, Hamburg, Germany). The alignment of the VR headset’s position and the calibration of the interocular distance between its two displays were adjusted for each participant to optimize visual acuity.

Before performing the main experimental task (introduced in the *Task and Setup* section), participants completed an initial assessment of the stimuli. In this phase, each animal was presented once for a duration of 3,000 ms at a fixed distance of 3.5 m (resulting in a total of 8 trials). After the offset of each animal, participants answered the following questions: (1) did you recognize the object? (“Yes”/“No”); (2) how threatening was the animal to you? (7-point Likert-type scale; min: “not at all”, max: “very much”); (3) how disgusting was the animal to you? (7-point Likert-type scale; min: “not at all”, max: “very much”); (4) how fast could the animal move towards you? (7-point Likert-type scale; min: “not at all”, max: “very fast”). Questions and answer options were displayed in the VR headset and participants verbally indicated their response that was recorded by the experimenter.

In a next step, to become familiar with the procedure of estimating distances using the VR controller, participants engaged in a practice run with each animal model (i.e., 8 trials in total). In these trials, the animal remained continuously visible at a random distance (uniformly distributed between 1.5 and 5.5 m) and participants had to place the distance marker at the closest visible part of the animal’s head.

In the main experimental trials (720 in total), the protocol was largely similar, but now the animals were presented for only 100 ms (see Figure 1a). The experimental trials were divided into 6 blocks, with each animal being shown 15 times per block. The sequence of animals within a block was randomized. The presentation of stimuli occurred independently of the participant’s cardiac phase, thus approximating a uniform distribution of stimulus onsets across the cardiac cycle. After each stimulus presentation, the entire scene was grayed out for 1,000 ms, which served as a backward mask and ensured that localization (in the afterwards empty room) was not simply based on the retinotopic percept or an afterimage of the scene with the animal (Coltheart, 1980; Raab, 1963). Participants undertook a minimum of five practice trials of the main task immediately before the first experimental block until they were comfortable with the procedure. The animal models used in these practice trials were distinct from those featured in the rest of the experiment.

Furthermore, in addition to the initial assessment of the stimuli, a shortened version was conducted prior to each experimental block to assess the participants’ recognition of the animals and the subjectively perceived level of threat *throughout* the experiment. This time, each animal was presented for a duration of 100 ms (at a random distance between 1.5 m and 5.5 m). Participants had to indicate whether they recognized the animal („Yes”/”No”) and rate the perceived level of threat posed by each animal (7-point Likert-type scale; min: “not at all”, max: “very much”).

After each block, the experiment was paused and participants could take a break and remove the VR headset. On average, participants completed the main part of the study, including breaks and the block-by-block stimulus assessments, in 65.0 minutes (*SD* = 9.7 minutes, range: 50.0–92.0 minutes). The practice section, along with the initial stimulus assessments, took an average of 6.4 minutes to complete (*SD* = 1.2 minutes; range: 3.9–10.6 minutes). Upon completion of all experimental trials, the participants again completed the shortened SSQ for a post-exposure measure of symptoms of cybersickness as well as the Slater-Usoh-Steed Questionnaire (SUS; Slater, 1999) to measure the level of presence in the virtual environment.

### ECG data preprocessing

We used Matlab R2019b (Mathworks, Natick, MA, USA) and the toolboxes EEGLAB v2019.1 (Delorme & Makeig, 2004) and HEPLAB v1.0.1 (Perakakis, 2019) for preprocessing the ECG data. To determine the onset of each cardiac cycle (i.e., detect the R peak), we first band-pass filtered the data (non-causal zero-phase FIR filter with hamming window of length 6601 samples, lower/upper passband edge: 0.50/40.00 Hz, transition bandwidth: 0.50 Hz, lower/upper –6 dB cutoff frequency: 0.25/40.25 Hz) and then applied functionalities of HEPLAB (*heplab_fastdetect.m*, based on De Carvalho et al., 2002). We applied visual inspection to identify noisy stretches of the ECG data affected by artifacts (e.g., high-frequency noise) which rendered the recognition of R peaks impossible (see *Data exclusion* section). For each trial, the timing of stimulus onset relative to the RR interval was then classified using both a circular and a binary approach to account respectively for the oscillatory and biphasic nature of cardiac activity (for an extensive description see: Al et al., 2020; Kunzendorf et al., 2019; Motyka et al., 2019). The circular approach relies on determining the relative position of the stimulus between one heartbeat and the next (taking values from 0 to 2π; Pewsey et al., 2013). Thus, it not only allows for consideration of the entire length of the RR interval but also normalizes for both inter- and intraindividual variability. In the binary approach, the cardiac cycle is segmented into systolic and diastolic phases with the use of an algorithm for the detection of the end of the T wave in the ECG (Vázquez-Seisdedos et al., 2011). Systole was defined as the interval from 50 ms after the R peak to the T wave end, while diastole was defined as the interval from 50 ms after the T wave end to 50 ms before the following R peak. This method accounts for both within- and between-subject variations in the duration of systole and diastole (in milliseconds) and facilitates the comparison of the obtained results with prior studies, which typically use the two-phase distinction. The binning procedure yielded on average 251.9 systolic and 315.0 diastolic trials per participant which reflects the difference in the average length of the respective cardiac phases (279.6 ms vs 363.9 ms).

### Analysis

We focused on the deviance between the reported position of the animal and its actual position (during the presentation) as the outcome variable of interest, which we term *localization error*. To conduct our analyses and interpretations, we broke down this error into two components (see Figure 1b):

#### Distance error

The difference in length between two vectors – one from the participant to the reported location of the stimulus animal and the other to its actual location. Negative distance errors indicate an underestimation of the distance to the animal. If threatening objects are perceived as closer than their actual distance (especially when presented during cardiac systole), we would expect a negative bias in the distance error for these specific animals (hypothesis 1), particularly in the systolic trials (hypothesis 2).

#### Angular error

The angle between the vector from the participant to the reported location of the stimulus animal and the vector to its actual location.

As we were interested in the perceived proximity of objects as a function of their threat level and the cardiac phase during which they are perceived, we focused our analyses on the distance error component (being the primary measure of interest).

To test hypothesis 1, we applied a paired, two-tailed *t* test on the group level to assess whether the average distance errors for threatening animals significantly differed from those for non-threatening animals. To assess the overall accuracy of the distance estimates (i.e., the presence of a global bias to over- or underestimate), we performed a one-sample, two-tailed *t* tests against zero, pooling the average distance errors across all experimental conditions.

In a next step (hypothesis 2), we extended the analyses to repeated measures analyses of variance (rmANOVA) models which, besides the binary main effect *threat* (with the two levels threatening and non-threatening animals) also included the predictor *cardiac phase* (with the two levels *systole* and *diastole*) as well as their interaction. This allowed us to corroborate whether the cardiac phase during which the animal was perceived had an effect on the distance error.

To assess whether non-significant results reflected null effects rather than experimental insensitivity, we calculated Bayes factors for the comparisons of interest (Dienes, 2014). The default JZS prior (r = 0.707), was used, given the absence of empirical evidence or a quantitative theoretical model that could inform prior specification (Rouder et al., 2012). The relative robustness of Bayes factors was further validated in analogous comparisons using different prior widths (narrow, r = 0.354, and wide, r = 1; van Doorn et al., 2021).

To increase the sensitivity of our analyses, we additionally applied linear mixed-effects models (as implemented in the statistical software package lme4; Bates et al., 2015), which account for single-trial data and allow us to model a random effects structure (e.g., random intercepts per participant to account for interindividual variations). In the first model, we treated the cardiac phase as a binary predictor for the distance error, contrasting cardiac systole and diastole (dummy-coded; diastole: 0). The binary fixed effect *threat* represented whether the stimulus animal in a given trial was threatening or not (dummy-coded; non-threatening: 0). To account for interindividual differences, we fitted random intercepts for individual participants. Furthermore, pilot data suggested that the distance error varies depending on the true distance at which the stimulus has been shown. As this effect might differ across individuals, we included the *actual distance* (blue line in Figure 1) as a continuous random effect in the model.

Finally, we adapted this model to incorporate the cardiac phase as a continuous, circular predictor, which was formalized as a combination of its sine and cosine components (Pewsey et al., 2013). This approach eliminates the need for an a priori definition of relevant time-windows and tests the hypothesis of a non-uniform distribution of the outcome variables (distance and angular error) across the interval between two R peaks.

As the specification of these models allows for many degrees of freedom, we had pre-registered the reported models before acquiring the data (https://osf.io/a7n9b/). All statistical analyses were conducted using the R statistical software (v4.0.1; R Core Team, 2021) and RStudio (RStudio Team, 2021). All data and code used for the analyses are publicly available (data: https://doi.org/10.17617/3.KJGEZQ, code: https://github.com/eioe/vrcc_analysis).

### Control analyses

We examined if the second component of the localization error, namely the (absolute) angular error, was modulated by the threat level of an animal or the cardiac phase. To achieve this, we employed the same 2×2 rmANOVA (threatening vs non-threatening animals; systolic vs diastolic presentation) that we used for modeling the distance error.

Additionally, we employed a similar analysis to assess whether precision, defined as the width of the individual participants’ error distributions, displayed variation between threatening and non-threatening animals, as well as between systole and diastole. To accomplish this, we computed the standard deviation of the distance errors and the angular errors across trials within a given condition for each participant. Subsequently, we used these values on the group level as the dependent variable, separately for both types of errors.

### Data exclusion

The following number of trials were excluded (based on pre-registered criteria): 130 trials with no distance estimation response within 5 seconds (from 28 participants); 112 trials with a noisy ECG signal precluding reliable determination of the R peak and the cardiac phases (from 13 participants); 7 trials with exceptionally large localization errors (>3 SD from the mean; from 7 participants); 388 trials with irregularly short or long systolic interval (>3 SD from the mean; from 41 participants). No subjects were excluded due to a deviation of their average distance error (>3 SD from the mean) or an insufficient number of trials (< 70% trials remaining after application of all trial rejection criteria). Overall, the average proportion of trials retained for analysis per individual was 98.0% (*SD* = 1.7%; range: 89.4–99.9%).

## Results

### Perceived distance to threatening and non-threatening stimuli (hypothesis 1)

Hypothesis 1, that observers perceive threatening animals as closer than non-threatening ones, was not supported. We found a significant difference between the distance errors (*t*(40) = 3.34, *p* = .002, Cohen’s *d* = 0.07). However, contrary to the hypothesis, the mean distance error was significantly higher (*M* = 0.78 cm, *SD* = 30.68 cm) for threatening than for non-threatening animals (*M* = −1.34 cm, *SD* = 31.38 cm). This indicates that, on average, distances to non-threatening animals were slightly under- and distances to threatening animals marginally over-estimated (Figure 2a). Notably, this effect was driven by the overestimations for a single animal (snake), and no clear pattern contrasting the threat conditions emerged (see Figure 2). Across all animals, the mean distance error (*M* = −0.28 cm, *SD* = 30.97, range: –90.66–75.38) was not significantly different from 0 cm as corroborated by a two-sided, one-sample *t* test (*t*(40) = −0.06, *p* = .95).

**Figure 2.**
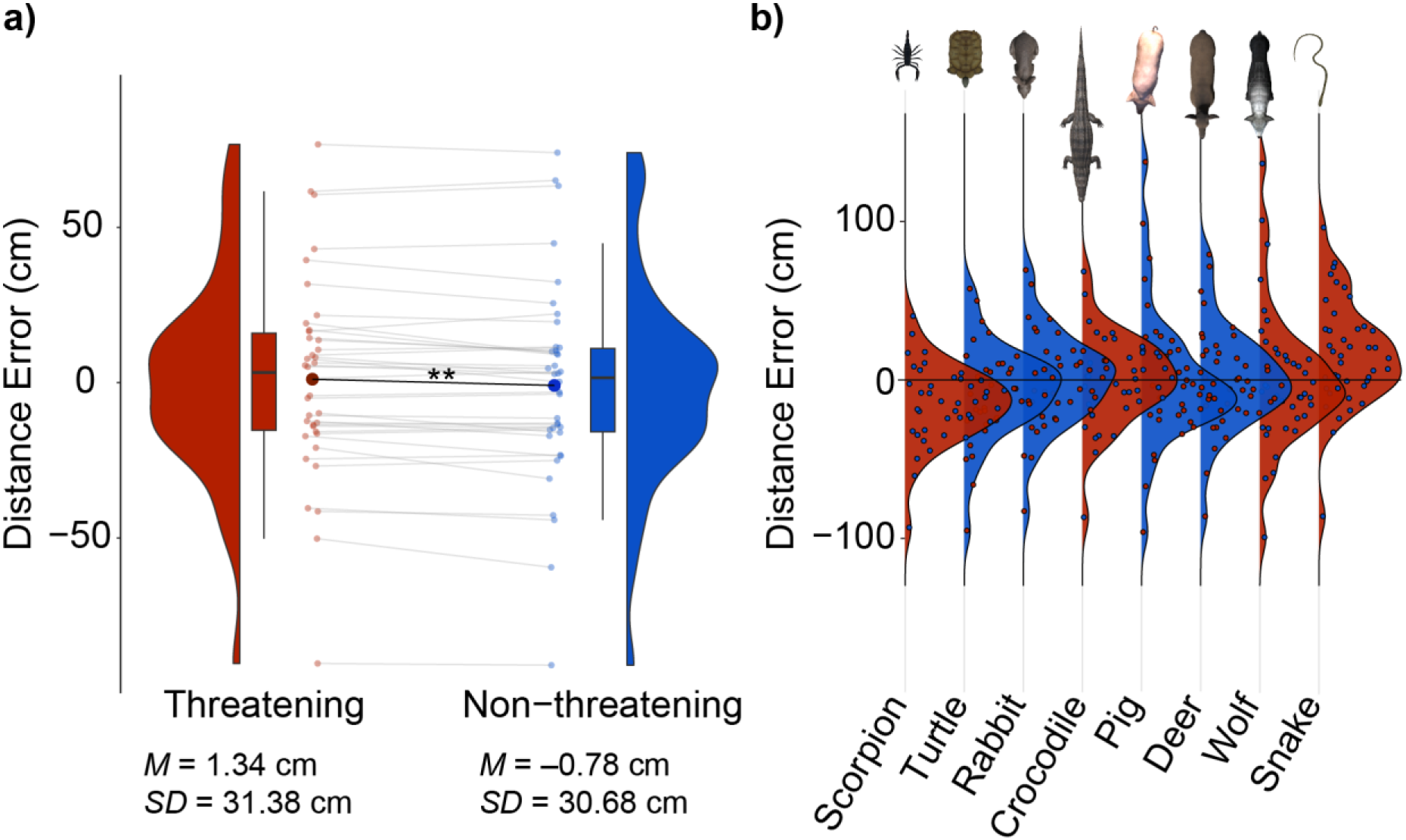
(**a**) The average distance error – that is the difference between the indicated and the actual distance – was significantly but only marginally (by 2.12 cm) lower for non-threatening than for threatening animals. Negative and positive values correspond to, in that order, underestimation and overestimation of the indicated distances. (**b**) Average distance errors for individual stimuli (red – threatening; blue – non-threatening animals) in increasing order. ** *p* < .01.

### Perceived distance to stimuli at different cardiac phases (hypothesis 2)

The second hypothesis, postulating a decreased perceived distance to threatening stimuli at earlier phases of the cardiac cycle (i.e., systole) relative to the later phases (i.e., diastole), was tested using both binary and circular approaches. A 2×2 rmANOVA with cardiac phase (systole/diastole) and threat (threatening/non-threatening animals) as binary factors and average distance error as the dependent variable confirmed the significant main effect of threat (*F*(1, 40) = 7.87, *p* = .008, η^2^*_G_* = .001) but both the main effect of cardiac phase (*F*(1, 40) = 0.11, *p* = .746, η^2^*_G_* < .0001) and its interaction with threat (*F*(1, 40) = 0.94, *p* = .337, η^2^*_G_* = .0001) were non-significant (Figure 3b). The only (Bonferroni-corrected) significant difference in the post-hoc comparisons was the difference between threatening and non-threatening animals at systole (in the opposite direction to the one hypothesized: *p_Bonferroni_* = .045; the result of an analogous comparison for diastole was not significant: *p_Bonferroni_* = .111).

**Figure 3.**
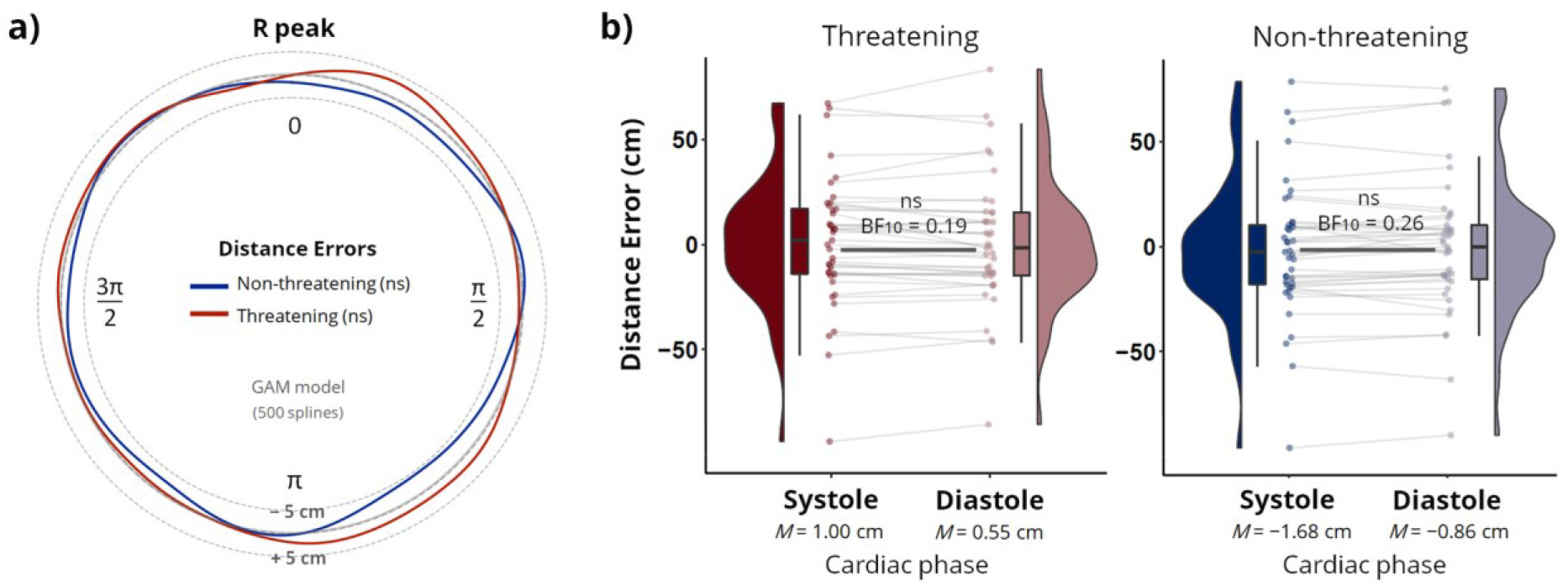
Circular and binary analysis of perceived distance to threatening and non-threatening animals relative to the heartbeat. (**a**) Non-linear smooths represent fluctuations in the average distance error to particular stimuli across the entire cardiac cycle (from R peak to R peak). The dotted lines indicate distance errors 5 cm below and above the true distance. We applied a generalized additive model (GAM): First, a regression model with 500 splines was used to estimate the cardiac phase-dependent variations in distance error separately for each participant and condition (using Python’s PyGAM library, version 0.8.0, with default terms; Servén et al., 2018). The individual non-linear smooths were then averaged to illustrate the mean for the entire sample. In sum, we did not find evidence for cardiac phase-dependent variations in the perceived distance to threatening and non-threatening animals. (**b**) The binary analysis – that segments the cardiac cycle into systole and diastole based on a T wave end detection algorithm – also did not indicate significant between-phase differences in distance error. The corresponding Bayes factor analysis yielded substantial evidence for the null hypothesis (*BF_10_* < 0.33, ns = non-significant).

A Bayesian analysis of the non-significant results regarding the cardiac phase, yielded substantial evidence for the null hypothesis (*BF*_10_ < ⅓) that distance perception does not differ between systole and diastole, both for threatening (*BF_10_ =* 0.19) and non-threatening animals (*BF_10_ =* 0.26; Figure 3b). This finding remained also with a narrower (r = 0.354; threatening: *BF_10_* = 0.34; non-threatening: *BF_10_* = 0.45) and a wider prior (r = 1: threatening: *BF_10_* = 0.14; non-threatening *BF_10_* = 0.19).

Analyses of cardiac phase differences in distance error for individual animals separately showed no significant results for any of the animals (all *p* values > 0.33; Figures S2, S3).

Also in the (more sensitive, pre-registered) linear mixed model with threat and cardiac phase both as binary predictors, only the main effect of threat was significant (*β* = 1.44, *SE* = 0.74, *t* = 1.97, *p* = .049), while the main effect of cardiac phase (*β* = −0.87, *SE* = 0.78, *t* = −1.11, *p* = .267), as well as their interaction (*β* = 1.18, *SE* = 1.10, *t* = 1.07, *p* = .286), were not significant. This was further confirmed by the linear mixed model in which we modeled the distance error as a function of the cardiac phase (at the moment of stimulus presentation) as a continuous, circular predictor (formalized as a combination of sine and cosine) and threat (binary). Also for this model, only the main effect of threat was significant (*β* = 2.16, *SE* = 0.49, *t* = 4.13, *p* < .001), while the circular factors regarding the cardiac phase as well as their interactions with threat were not significant predictors of the distance error (sine: *β* = −0.71, *SE* = 0.49, *t* = −1.45, *p* = .147; cosine: *β* = 0.54, *SE* = 0.49, *t* = 1.11, *p* = .269; sine * threat: *β* = 0.99, *SE* = 0.69, *t* = 1.43, *p* = .154; cosine * threat: *β* = −0.65, *SE* = 0.69, *t* = −0.94, *p* =.347).

### Control analyses

The (absolut) angular error did not vary as a function of an animal’s threat level or the cardiac phase during which it was perceived. A 2×2 rmANOVA did not reveal a significant main effects of threat (*F*(1, 40) = 0.125, *p* = .726) or cardiac phase (*F*(1, 40) = 0.007, *p* = .935) nor a significant interaction (*F*(1, 40) = 2.236, *p* = .143).

Assessing the precision (i.e., the width of the error distributions as summarized by the standard deviation) between threatening and non-threatening animals and between systole and diastole, we found a similar pattern as for the mean of the distance error. We observed a significant main effect of threat for the precision of the distance error (*F*(1, 40) = 4.33, *p* = .044) as well as for the angular error (*F*(1, 40) = 16.45, *p* < .001), but not for the main effect of cardiac phase (distance error: *F*(1, 40) = 0.002, *p* = .961; angular error: *F*(1, 40) = 0.24, *p* =.624) or their interaction (distance error: *F*(1, 40) = 0.22, *p* = .645; angular error: *F*(1, 40) = 0.34, *p* =.562). The precision was higher (i.e., the average standard deviation was lower) for non-threatening as compared to threatening animals, in terms of distance errors (*M*_threat_ = 31.72 cm, *SD*_threat_ = 12.18, *M*_non-threat_ = cm, *SD*_non-threat_ = 11.65) as well as angular errors (*M*_threat_ = 2.18°, SD_threat_ = 0.45, *M*_non-threat_ = 2.01°, *SD*_non-threat_ = 0.41).

### Subjective ratings of the stimuli and questionnaire results

The initial stimuli assessment (during which the animals were shown for three seconds) confirmed the results from the online study. In the recognition task, participants showed no considerable difficulty in identifying briefly (100 ms) presented animals (non-identifications or no answers were registered in only 93 out of 1968 cases). Animals classified as “threatening” based on the online study were evaluated as significantly more threatening (*M* = 4.57, *SD* = 1.63) than animals originally classified as “non-threatening” (*M* = 1.31, *SD* = 0.38, *t* = 13.6, *p* < .001; see Figure S4 for individual animal ratings). Threatening animals were further rated as significantly more disgusting (*M* = 2.74, *SD* = 1.48) than non-threatening ones (*M* = 1.34, *SD* = 0.43, *t* = 6.93, *p* < .001; Figure S5) as well as significantly faster (*M* = 5.06, *SD* = 0.97) than the non-threatening animals (*M* = 3.52, *SD* = 0.87, *t*(40) = 10.72, *p* < .001; Figure S6).

The average rating score from the six threat ratings preceding the experimental blocks (100 ms presentation time per animal) was also higher for threatening (*M* = 4.26, *SD* = 1.81) than for non-threatening animals (*M* = 1.28, *SD* = 0.42, *t* = 11.47, *p* < .001). To assess whether animals became less threatening over the course of the experiment, we examined the block-by-block changes in the subjective ratings of threatening animals before each experimental block. We observed significant differences (*F*(2.7, 108.1) = 4.43, *p* = .007, ε = 0.54, η^2^*_G_* = .006; GG-corrected; Figure S7) between blocks 1 (*M* = 4.49) and 6 (*M* = 4.05, *p_Bonferroni_* = .001) as well as blocks 1 and 5 (*M* = 4.11, *p_Bonferroni_* = .009). Yet, absolut decreases in threat ratings were moderate and ratings for threatening animals (*M* = 4.05, *SD* = 1.85) were still significantly higher (*t*(40) = −10.26, *p* < .001) than for non-threatening animals (*M* = 1.31, *SD* = 0.53) in the final rating round.

The initial and mid-experimental threat ratings for threatening animals were highly correlated. Comparing the average across all mid-experimental ratings per animal with the corresponding initial ratings yielded Spearman correlation coefficients ranging from 0.58 to 0.76. Therefore, in the subsequent analyses that include individual threat ratings per animal, we refer to the average from all 7 ratings (1 rating with long, 3,000 ms, presentation times at the beginning of the experiment + 6 ratings with short, 100 ms, presentation times before the beginning of each block). As for the post-experimental questionnaires, the level of self-reported presence in the virtual environment was at an average level (*M* = 3.80, *SD* = 1.08, range: 1.67–5.83, on the 7-point SUS questionnaire scale). After exposure to the VR environment, we observed a small significant increase (*t* = 2.80, *p* = .008) in self-reported symptoms of cybersickness (*M* = 1.45, *SD* = 0.40) as compared to their baseline level (*M* = 1.22, *SD* = 0.30). Notably, the procedure did not evoke severe symptoms of cybersickness in any of the participants (range of the mean scores after the experiment: 1–2.67 on the 7-point SSQ scale).

### Perceived distance to threatening stimuli as a function of subjective threat, disgust, and speed ratings

To explore whether perceived distance to stimuli decreased with subjective threat, we analyzed correlations between participant- and animal-specific distance errors and subjective feelings of threat evoked by specific animals. Given the minimal variability in threat ratings for non-threatening animals (floor-level scores), they were dropped from the analysis. In short, we did not find significant associations between distance error and threat ratings for any of the threatening animals (crocodile *r_s_* = .02, *p* = .898; wolf *r_s_* = −.23, *p* = .143; snake *r_s_* = −.08, *p* = .616; scorpio *r_s_* = −.02, *p* =.908; Figure S8). An analogous analysis was performed to explore whether perceived distance increased with subjective disgust towards particular animals – sampled only once at the start of the experiment (see Cole et al., 2013). Also here, only threatening animals were analyzed due to insufficient variability of disgust ratings for the non-threatening animals. Again, there were no significant relationships between distance error and subjective feelings of disgust (crocodile *r_s_* = −.11, *p* = .500; wolf *r_s_* = −.04, *p* = .811; snake *r_s_* = −.20, *p* = .218; scorpio *r_s_* = −.13, *p* = .425). Finally, the ratings of expected movement speed were also not significantly correlated with distance error (both for threatening: crocodile *r_s_* = .04, *p* = .797; wolf *r_s_* = .09, *p* = .599; snake *r_s_* = −.01, *p* = .958; scorpio *r_s_* = .20, *p* = .221; and non-threatening animals: pig *r_s_* = −.16, *p* = .318; deer *r_s_* = −.12, *p* = .443; turtle *r_s_* = .08, *p* = .616; rabbit *r_s_* = .123, *p* = .444).

### Perceived distance to stimuli as a function of anxiety levels

We also explored whether there was a significant association between participants’ anxiety levels, as measured by the State-Trait Anxiety Inventory (STAI-S and STAI-T), and perceived distances to presented stimuli. No significant correlation was found between state anxiety and distance error averaged across all animals (*r_s_* = −.17, *p* = .287). This also held when tested for each animal individually (range *r_s_*: −.05 to −.18; *p*: .256 to .769). By contrast, we observed a significant negative correlation between trait anxiety and averaged distance error (*r_s_* = −.36, *p* = .020; Figure S9). The propensity to indicate stimuli as physically closer with increasing levels of trait anxiety appeared to be fairly consistent across different animals both threatening (crocodile *r_s_* = −.27, *p* = .083; wolf *r_s_* = −.36, *p* = .020; snake *r_s_* = −.27, *p* = .092; scorpio *r_s_* = −.34, *p* = .029) and non-threatening ones ( pig *r_s_* = −.33, *p* = .036; deer *r_s_* = −.34, *p* = .032; turtle *r_s_* = −.34, *p* = .030; rabbit *r_s_* = −.33, *p* = .036).

## Discussion

In this study, we investigated the effects of cardiac signals and subjective feelings on the perception of threatening and non-threatening animals in a naturalistic experiment using immersive virtual reality. Participants indicated the distance at which they perceived the briefly displayed animals while their cardiac activity was recorded. Neither of the two pre-registered hypotheses (https://osf.io/a7n9b/) was confirmed as we did not find that threatening animals are perceived as closer than non-threatening ones (Cole et al., 2013) and during earlier (i.e., systole) compared to later (i.e., diastole) phases of the cardiac cycle. On average, localization was precise and rather influenced by other (e.g., physical) stimulus characteristics than the threat level. Results from Bayesian analyses additionally provide substantial evidence for the absence of a cardiac phase bias in our data. Notably, our experiment adopted a more naturalistic approach, implementing an immersive task with a behavioral outcome measure, than classical experiments which found such effects (e.g., Azevedo et al., 2017; Cole et al., 2013; Garfinkel et al., 2014). In the following, the results and possible implications will be discussed per hypothesis.

Regarding our hypothesis 2, we did not observe that the perceived distance of threatening objects varied over the cardiac cycle. This outcome deviates from previous research that reported heightened processing of fear stimuli during cardiac systole. For instance, fearful faces, when presented during systole compared to diastole, elicit greater activity in the amygdala (Garfinkel et al., 2014) and are rated as more intense (Garfinkel et al., 2014; Leganes-Fonteneau et al., 2021; but see also Pfeifer et al., 2017). At the systolic phase, fearful faces capture visual attention more strongly, specifically at their low spatial frequencies (Azevedo et al., 2018) and are more easily detected in an attentional blink paradigm with backward masking (Garfinkel et al., 2014), but not in a visual search task (Leganes-Fonteneau et al., 2021). Besides faces, the threat level of handheld objects was shown to vary over the cardiac cycle, leading to an increased stereotype-driven misidentification of harmless objects as weapons during systole (Azevedo et al., 2017). While evidence supports a cardiac phase bias in the perception of visual threats, it remains unclear how well it generalizes across experimental paradigms (i.e., stimuli and designs). Here, we tested whether we would find effects of the cardiac cycle in a more naturalistic and immersive setting than those used in previous studies. To test this hypothesis, we probed the effects of the cardiac cycle on a concrete behavior (i.e., indicating the location at which a threatening animal was perceived) in immersive, stereoscopic virtual reality. Despite participants’ ability to perform this task (on average) with high precision and our dependent variable (i.e., the two distance errors) being sensitive to even minor deviations in the visual nature of the stimuli, we found no evidence for a modulation by the cardiac phase. In fact, Bayesian analyses provided substantial support for the null hypothesis, suggesting that the distance errors in systole and diastole were equal.

These null findings allow for (at least) two interpretations. First, there is no or only a negligible impact of the cardiac phase on distance estimation and related behaviors in naturalistic settings. Cardiac cycle effects which are measurable in more artificial and abstract setups and behavioral tasks (e.g., button presses), might be overruled by stronger effects once the situation becomes more lifelike and the queried behavior is more natural (e.g., indicating a concrete location in 3D space). To substantiate this interpretation, additional studies investigating cardiac cycle effects in setups with a high level of naturalism are needed. Ideally, such studies should implement different paradigms and behavioral outcome measures that are comparable in the level of naturalism to the task we used here, to clarify whether the absence of a cardiac cycle effect in our data was due to the increased level of naturalism. A second explanation could be that our novel experimental paradigm (presenting participants with naturalistic 3D renderings of more and less dangerous animals) was not suitable to detect the hypothesized effects. However, our rationale was based on studies that found cardiac cycle effects to be particularly strong for fear- and threat-related stimuli (Azevedo et al., 2017, 2018; Garfinkel et al., 2014; Leganes-Fonteneau et al., 2021). We set out to translate these previous results in a setup and task of increased naturalism (using immersive, stereoscopic presentation and distance estimation of photorealistic stimuli). At the same time, we maintained a rigorous level of experimental control by strictly timing the visibility of animals at well-defined locations and incorporating a high number of trials.

Concerning hypothesis 1, we did not find evidence for an overall bias to underestimate the distance of threatening compared to non-threatening stimuli. Contrary to our hypothesis, non-threatening animals were on average perceived as significantly closer than threatening ones. This effect, however, was driven by a single stimulus animal, the snake, whose distance was strongly and consistently overestimated by most subjects. It is worth noting that also the animal that was perceived as closest, the scorpion, belonged to the threatening category. In this instance, we found that the average degree of underestimation corresponded to the length of the scorpion’s pincers, which protruded beyond its forehead – the reference point for determining the animal’s objective position. The snake, on the other hand, had its head raised and extended forward. It is plausible that due to the brief presentation time, participants have memorized the position of the coiled body on the ground more readily than that of the elevated head. Furthermore, we observed similar patterns in the angular errors (Figure S10): For instance, the pig was presented with a slight (10°) rotation to the right to enhance its recognizability. This resulted in a consistent rightward bias in the position estimations. Conversely, a leftward bias was observed for the snake, whose body was situated primarily to the left of its head from the participants’ viewpoint. Hence, localization (and therefore distance processing) seems to have been less influenced by the animal’s threat level than by other stimulus features such as body shape and orientation.

The lack of observed influence of the animals’ threat level on distance estimates in our study might stem from our choice of stimuli or the experimental task. We aimed for a high degree of naturalism while maximizing the comparability between threatening and non-threatening stimuli. Therefore, we chose animals for both categories which approximately matched in size and other characteristics, to minimize the probability that such confounds would drive differential effects between the groups. We instructed and trained participants to point at the animal’s foremost point of the head (typically the “nose”) to ensure consistency in the measure (i.e., establish which point demarcated the animal’s actual position). Noteworthy, on average, participants’ localization estimates were remarkably precise (with a mean distance error of 0.28 cm). However, we still observed the aforementioned distortions, which we ascribe to each animal’s unique physical characteristics (theoretically orthogonal to their threat level). Perhaps threat-driven effects might have become apparent with stimuli lacking these specific physical characteristics. However, increasing the lifelikeness of stimuli in more naturalistic studies comes at the sacrifice of full controllability, which is easier to achieve for more simplistic or abstract stimuli. Experimental manipulations of interest (here: threat-level and cardiac phase) may become confounded or counteracted by conceptually irrelevant features (e.g., shape, color, speed) in naturalistic designs.

Importantly, this does not contradict the assertion that VR enables a high level of experimental control. While alternative experimental approaches (such as employing real animals as stimuli) render a certain level of control impossible, VR studies offer the potential for complete control over the visual environment. Yet, researchers may willingly trade some aspects of this control for other advantages, such as more naturalistic stimuli or conditions. Here, we decided to keep the animals as lifelike as possible, knowing that this constrains the interpretability of any comparison between the two groups. Despite our striving to match them in terms of physical size, threatening and non-threatening animals differed also on other dimensions than their threat level (e.g., feelings of disgust, speed, colors) which makes it difficult to determine which feature was driving a potential difference between the two groups.

Therefore, in addition to contrasting threatening and non-threatening animals, we examined the relationship between individual participants’ distance estimates and their subjective threat ratings – separately for each animal. This approach aimed to determine if those who felt more threatened by an animal also consistently rated its proximity differently (Cole et al., 2013). Such a within-animal analysis is less affected by the differences in physical characteristics between the animals. Yet, also in this analysis, we did not find evidence for an association between distance estimates and subjective threat levels (neither when using pre-nor mid-experimental threat ratings; see Cole et al., 2013 for a similar approach but divergent results). Overall, the results of the present study challenge the notion that feelings of threat reduce visually perceived distance to the feared objects. It has been argued that such a bias of the perceptual system may facilitate adaptive responses in dangerous situations (e.g., faster fight/flight response; Balcetis & Cole, 2014; Cole et al., 2013; de Carvalho, 2022). Notably, the studies supporting this claim were mostly based on verbal estimations of the distance (e.g., in inches) to threatening objects (such as a living tarantula; Cole et al., 2013; a person described as aggressive; Cole et al., 2013; or a pain-triggering button; Tabor et al., 2015), which constitutes a difference to our experiment. We strived for an experimental operationalization which minimizes the need for cognitive transformations (e.g., into an explicit metric), to be able to assess the influence of threat on immediate distance perception, location representation, and the resulting behavioral performance. Such less abstract, perception-related measures produce relatively more accurate and less bias-susceptible estimates compared to verbal ones (Andre & Rogers, 2006; Etchemendy et al., 2018; Kunz et al., 2009). The appointed distinction between judgment- and behavior-related reports is embedded in the broader discussion on whether the effects observed in studies using more abstract measures reflect true shifts in perceptual experience or rather non-perceptually driven changes in cognitive judgments (Firestone & Scholl, 2016, 2017; Schnall, 2017; Witt, 2017). This might explain the discrepancy between previous reports about distance estimates which were biased by the threat of the stimulus and our null findings. Switching from a (more abstract) verbal report task to a (mostly perception-driven) behavioral pointing task might have eliminated this bias. If, however, the underestimation of distances towards threats is rooted in its evolutionary benefits, this should be particularly reflected in concrete behavior, not only in more abstract cognitive or verbalized representations.

Lastly, in an exploratory analysis, we observed that persons with higher levels of trait anxiety reported smaller distances towards all animals. Supplementary figure S9 suggests that this observation is mostly driven by participants with high STAI-T values underestimating the actual distance to the animals (rather than persons with small STAI-T values overestimating the distances). However, such an interpretation in absolute terms is speculative. Among all personality traits, anxiety appears particularly potent in predicting attentional and perceptual functioning. For instance, trait anxiety has been linked to increased reliance on priors in perceptual decisions (Kraus et al., 2021), increased attentional bias toward threats (MacLeod & Mathews, 1988; Okon-Singer, 2018), increased scanning of the virtual environment in response to threats (Yilmaz Balban et al., 2021), and an increased extent of peripersonal space surrounding the face (Sambo & Iannetti, 2013). To our knowledge, our findings are the first to indicate an association of trait anxiety and a behaviorally reported reduction in the perceived distance to visual objects.

### Limitations

Methodological factors could have contributed to the absence of cardiac signaling effects in our study. One cause could be insufficient power due to too few trials or participants. Yet, our trial number (ca. 29k in total and 720 per participant) and sample size (n = 41) are higher than those of previous studies which reported such effects (e.g., Azevedo et al., 2017; Garfinkel et al., 2014). Furthermore, the results of Bayesian analyses indicate substantial support for the absence of cardiac phase bias in our data, rather than inconclusive evidence for distinguishing between the alternative explanations. Another limitation could be that our stimulation insufficiently induced a sense of threat. Importantly, our stimuli were carefully selected based on ratings collected in a separate sample, and the threat ratings by our actual participants also clearly differentiated between sets of animals classified as threatening and non-threatening. Moreover, menacing animals, particularly evolutionary threats like snakes and spiders (Fang et al., 2016; Öhman, 2009; Ruiz-Padial et al., 2005; Shibasaki & Kawai, 2009), share the same characteristic of rapid and automatic processing with fearful faces (Anderson et al., 2003; Kiss & Eimer, 2008; Pegna et al., 2008; Pessoa et al., 2005) for which cardiac phase effects have been consistently observed. Finally, cardiac phase biases were also found for non-facial objects that were in general less realistic than ours (Azevedo et al. 2017).

Moreover, 3D displays such as VR head-mounted displays (HMDs) necessarily involve a mismatch between focus cues: As the displays remain at a fixed distance from the eyes, focusing on the distance of the displays (accommodation) typically does not match the eye rotations to fixate the 3D object at its distance in the virtual scene (vergence). Such “vergence–accommodation conflicts” can cause eye strain and may contribute to an underestimation of egocentric distances in VR (Feldstein et al., 2020; Renner et al., 2013). This underestimation seems to become less pronounced with technological advances of VR HMDs such as higher field-of-view and visual resolution (Kelly, 2022) and can be alleviated with pictorial depth cues such as shadows or textures in the virtual environments (Ahn et al., 2020; Feldstein et al., 2020; Renner et al., 2013). In our study, this appeared to be the case, as we observed no overall bias in distance perception (with a mean distance error < 0.5 cm). Yet, with the outcome measure consistently proving its sensitivity to the physical characteristics of the stimuli, we argue that it was well-suited to reflect also other potential influences on distance estimation – for example, related to threat or the cardiac phase. On top of that, the use of a naturalistic 3D scenario in immersive VR should have even boosted the overall persuasiveness of threat in our study compared to classical experiments that typically use de-contextualized stimuli presented on 2D screens. For example, human escape decisions to animals and objects with different threat levels were successfully tested in VR (Sporrer et al., 2023). It was a core objective of our study to test the cardiac phase bias for visual threat perception in a more naturalistic scenario and using a situation-embedded behavioral measure. Even though it cannot be ruled out that other types of stimuli (e.g., fear-related expressions or individuals) would yield different results, under the present circumstances the cardiac cycle did not bias visual perception as measured by estimated distance.

In conclusion, our findings suggest that under immersive VR conditions, naturalistic and highly-detailed exteroceptive information (i.e., stimulus characteristics) determine visual distance estimation (i.e, the perceived proximity to threatening animals) to an extent that it overrides the potential effects of subjective feelings of threat and interoceptive signals from the heart.

## Acknowledgements

We thank Eva Kozáková, Stella Kunzendorf, Marta Paź, Petra Skalníková, Kristýna Křížová, and Luca Schulze-Buschoff for their helpful comments during the design preparation phase.

