## Supplementary Material for "No cardiac phase bias for threat perception under naturalistic conditions in immersive virtual reality"

* These authors contributed equally.

[1] Max Planck Institute for Human Cognitive and Brain Sciences, Neurology, Leipzig, DE
[2] Humboldt-Universität zu Berlin, Faculty of Philosophy, Berlin School of Mind and Brain, Berlin, DE
[3] Polish Academy of Sciences, Institute of Psychology, Warsaw, PL
[4] University of Warsaw, Faculty of Psychology, Warsaw, PL
[5] CausalPython.io
[6] Charles University, Third Faculty of Medicine, Prague, CZ
[7] National Institute of Mental Health, Czechia, Center for Virtual Reality Research in Mental Health and Neuroscience, Klecany, CZ
[8] Charles University, Third Faculty of Medicine, Prague, CZ

Corresponding authors:
Felix Klotzsche,
Paweł Motyka,

**Overview:**

Figure S1. Average threat ratings from an online pre-study
Figure S2. Between-phase differences in distance error for individual animals.
Figure S3. Distance errors in relation to true distances.
Figure S4. Average threat ratings from the main experiment.
Figure S5. Average disgust ratings from the main experiment.
Figure S6. Average ratings of expected movement speed from the main experiment.
Figure S7. Block-by-block changes in average threat ratings of threatening animals.
Figure S8. Associations between subjective threat ratings and distance errors.
Figure S9. Associations between anxiety levels and distance errors.
Figure S10. Angular error distributions per animal.

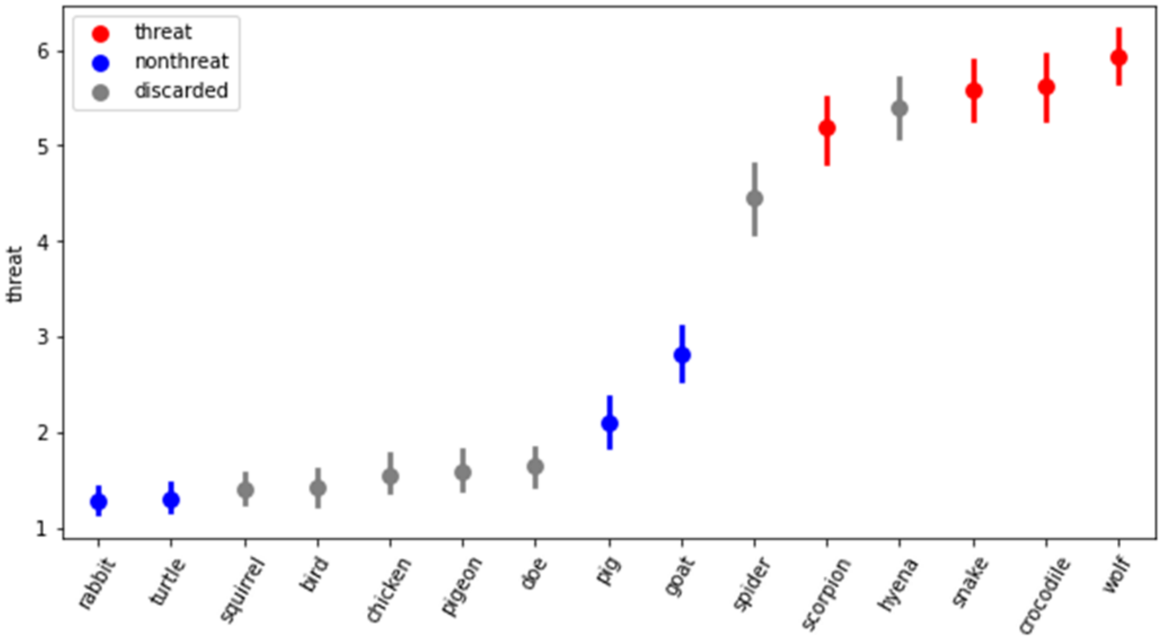


Figure S1. Average threat ratings from an online pre-study, in which an independent set of participants (n = 94) rated pictures of fourteen 3D animal models (error bars indicate 95% confidence intervals).

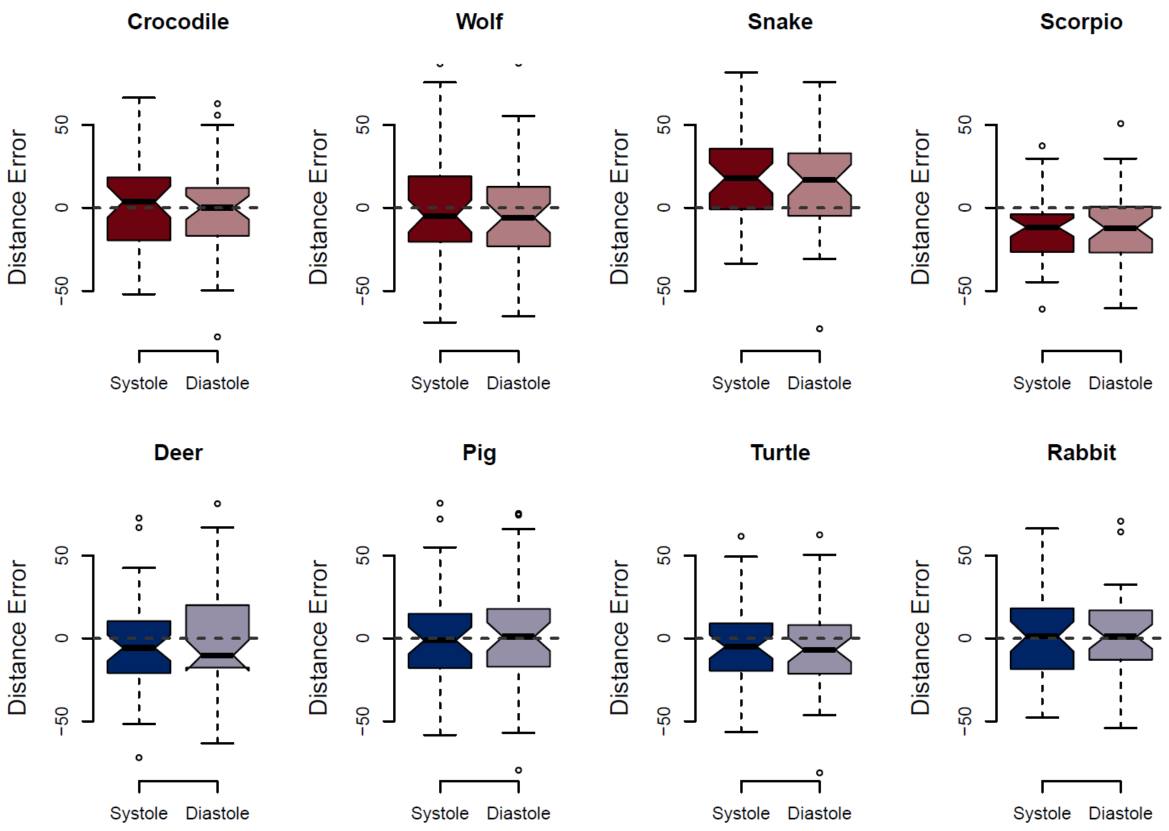


Figure S2. Between-phase differences in distance error for individual animals (all p values > 0.33).

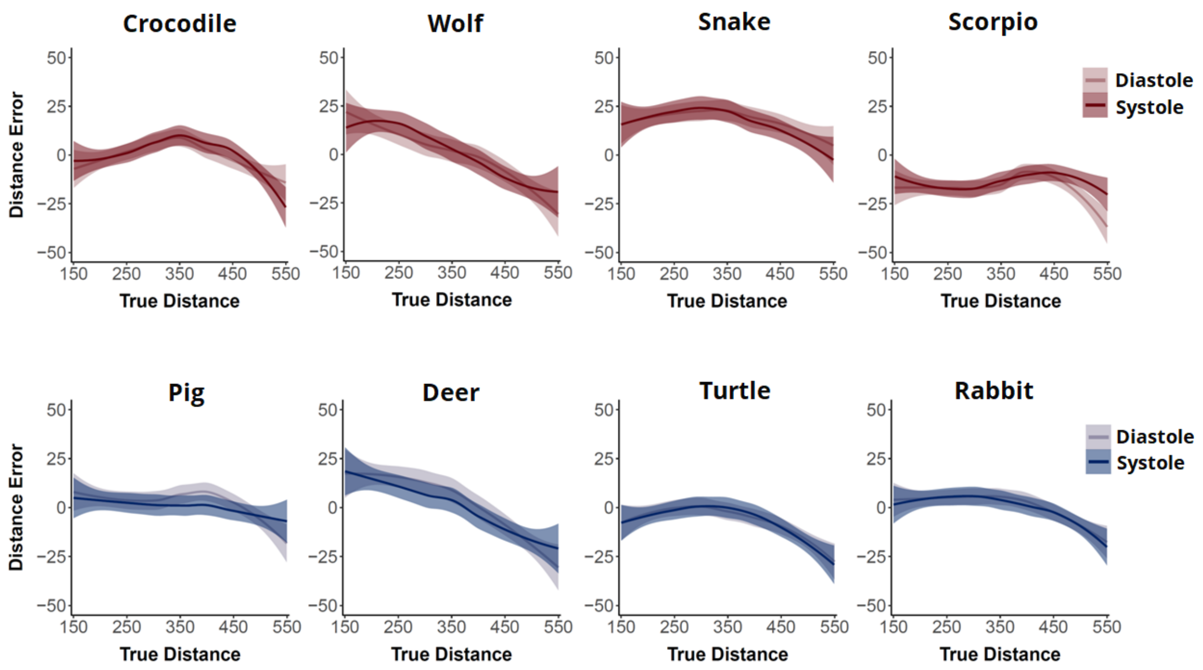


Figure S3. Distance errors in relation to true distances plotted based on raw data (i.e., without averaging at the subject's level) using the loess local regression fitting (coloured bands indicate 95% confidence intervals).

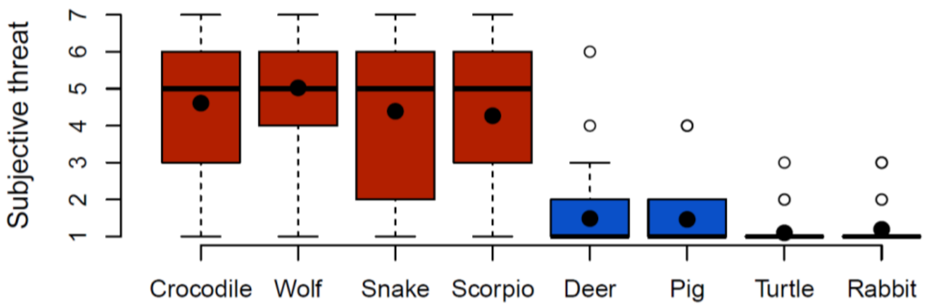


Figure S4. Average threat ratings from the main experiment (thick line: median; thick dot: mean).

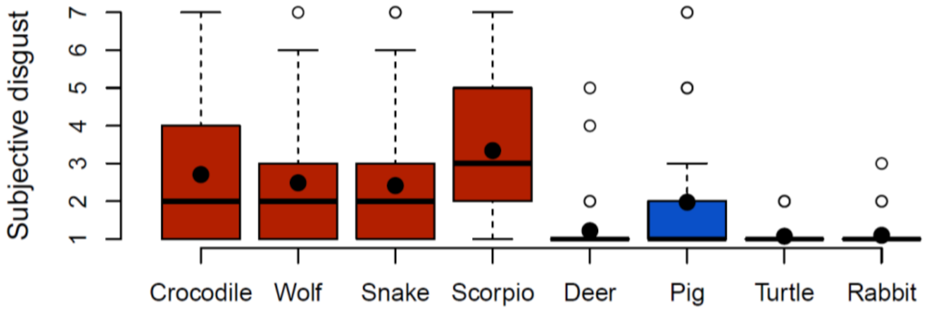


Figure S5. Average disgust ratings from the main experiment.

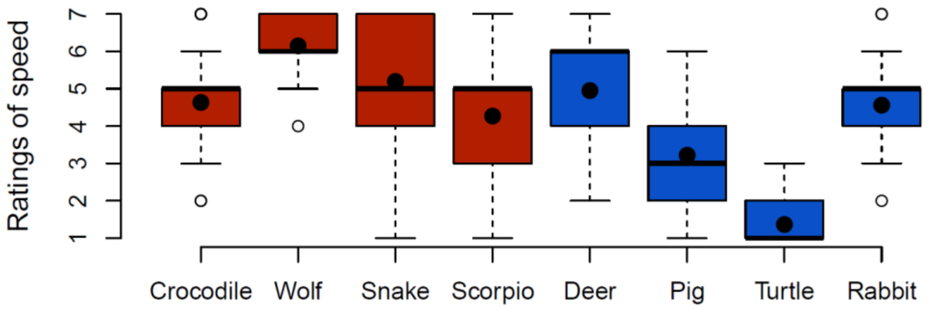


Figure S6. Average ratings of expected movement speed from the main experiment.

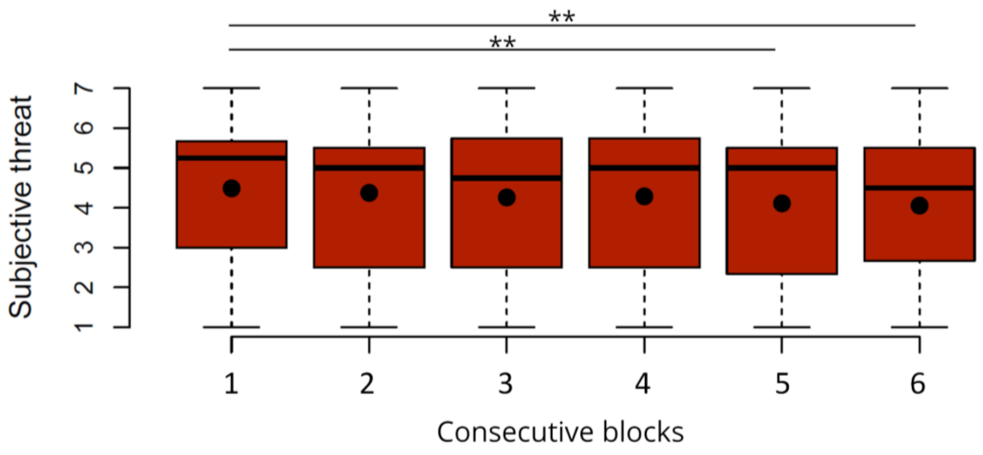


Figure S7.  Block-by-block changes in average threat ratings of threatening animals.

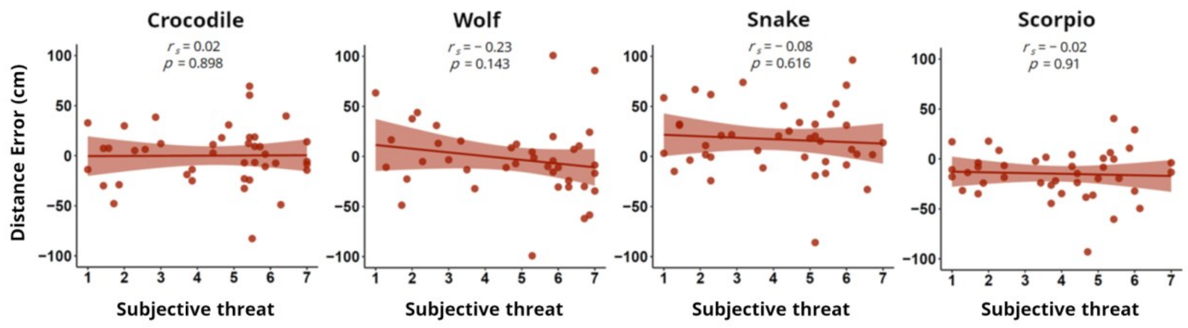


Figure S8. Associations between subjective threat ratings (x-axis) and distance errors (y-axis). Given the minimal variability in threat ratings for non-threatening animals (floor-level scores), they were dropped from the analysis.

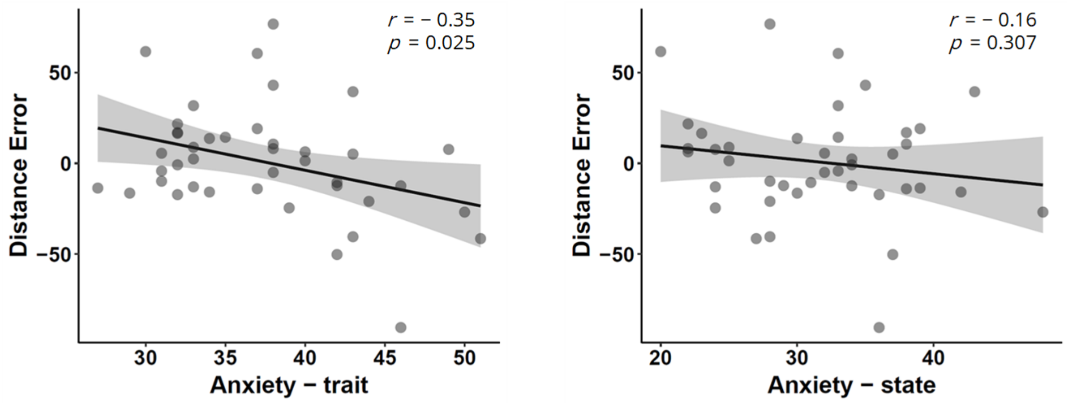


Figure S9.  Associations between anxiety levels (x-axis) and distance errors (y-axis).

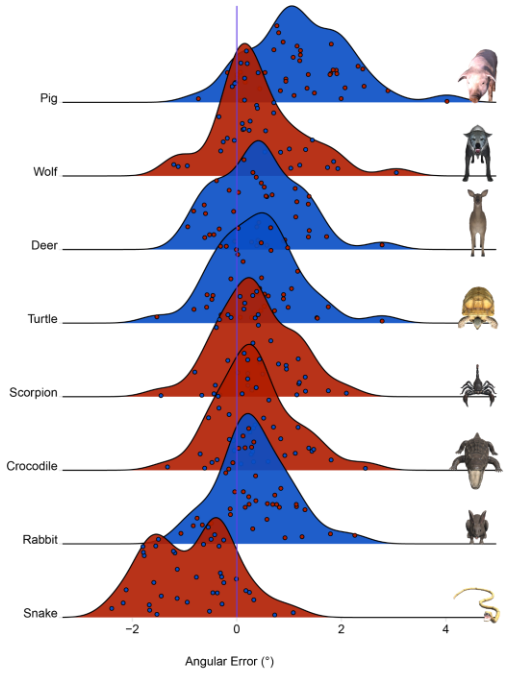


Figure S10. Angular error distributions per animal.
